# Cell cycle oscillations driven by two interlinked bistable switches

**DOI:** 10.1101/2023.01.26.525632

**Authors:** Pedro Parra-Rivas, Daniel Ruiz-Reynés, Lendert Gelens

## Abstract

Regular transitions between interphase and mitosis during the cell cycle are driven by changes in the activity of the enzymatic protein complex cyclin B with cyclin-dependent kinase 1 (Cdk1). At the most basic level, this cell cycle oscillator is driven by negative feedback: active cyclin B Cdk1 activates the Anaphase-Promoting Complex - Cyclosome, which triggers the degradation of cyclin B. Such cell cycle oscillations occur fast and periodically in the early embryos of the frog *Xenopus laevis*, where several positive feedback loops leading to bistable switches in parts of the regulatory network have been experimentally identified. Here, we build cell cycle oscillator models to show how single and multiple bistable switches in parts of the underlying regulatory network change the properties of the oscillations and how they can confer robustness to the oscillator. We present a detailed bifurcation analysis of these models.

## Introduction

Biological systems are often characterized by periodic behavior, ranging from the circadian clock, to the cell division cycle, and our heart rate. These biological rhythms are all generated by biochemical clock-like oscillators Winfree (2011); Goldbeter and Berridge (1996); Novák and Tyson (2008); Beta and Kruse (2017); Ferrell et al. (2011); Mofatteh et al. (2021). In the case of our heart rate, the sino-atrial node generates 50 to 150 action potentials per minute, depending on the oxygen demands of the body. Similarly, the period of the cell division cycle ranges from 10-30 minutes in rapidly cleaving early embryos to about 10-30 hours in dividing somatic cells. However, despite the large variation in the period of these oscillations, the amplitude remains reasonably unchanged. This has been attributed to the presence of positive feedback in the regulatory network of the biological oscillator Tsai et al. (2008). While any (biological) oscillator requires negative feedback to reset the system after each cycle Thomas (1981), positive feedback has been shown to be important to generate robust, large-amplitude oscillations with tunable frequency Brandman et al. (2005); Tsai et al. (2008). At the core of such *relaxation oscillations* is the fact that positive feedback can generate bistability in part of the regulatory network. Additional negative feedback can then drive oscillations along both branches of the bistable response curve. As a result, such relaxation oscillators exist over a wider range of parameters and allow the system to regulate its oscillation frequency without compromising its amplitude. Without a built-in bistable switch, changes in the frequency would strongly correlate with changes in the amplitude.

In the early embryo of the frog *Xenopus laevis*, it has been shown experimentally and theoretically that cell cycle oscillations are driven by a positive-plus-negative feedback network. There exists a (time-delayed) negative feedback loop between active enzymatic protein complexes of cyclin B with cyclin-dependent kinase 1 (Cdk1) and the Anaphase-Promoting Complex - Cyclosome (APC/C), which triggers the degradation of cyclin B Yang and Ferrell (2013); Rombouts et al. (2018). Positive feedback exists in the regulation of the activity of the Cdk1 complexes by the enzymes Wee1, Myt1 and Cdc25, which leads to bistability of Cdk1 activity in function of cyclin B concentration Pomerening et al. (2003); Sha et al. (2003) [see Figure 1(a),(c)]. However, measurements also show that the bistability in the dose-response curve of cyclin B vs. active cyclin B - Cdk1 is essentially abolished after the first cell cycle Tsai et al. (2014). Despite this loss of bistability, cell cycle oscillations persist and at a faster pace. Interestingly, various new experiments point towards the existence of a second bistable switch in the system, one where APC/C gets activated in a bistable manner by active Cdk1 Mochida et al. (2016); Rata et al. (2018); Kamenz et al. (2021) [see Figure 1(b)]. This bistable switch is related to the feedback loops involving Greatwall (Gwl), ENSA-Arpp19, and PP2A:B55 [see Figure 1(d)]. This triggers the question of what the advantage might be of interlinking bistable switches to generate cell cycle oscillations when combined with negative feedback? We recently found that having multiple bistable switches can increase the parameter range of oscillations, while also allowing to switch between regions of qualitatively different oscillations (in terms of period and amplitude) De Boeck et al. (2021). Having multiple positive feedback loops (which can lead to bistability) has been shown to allow for improving speed and resistance to noise Brandman et al. (2005), which could also indicate a potential advantage of interlinked bistable switches. Moreover, the cell cycle has also been studied as a chain of interlinked bistable switches, which can increase the robustness of cell cycle transitions Hutter et al. (2017); Novák and Tyson (2021). However, as far as we know, no study has looked in detail at how multiple bistable switches affect the dynamic properties of a resulting biological oscillator.

**Figure 1.**
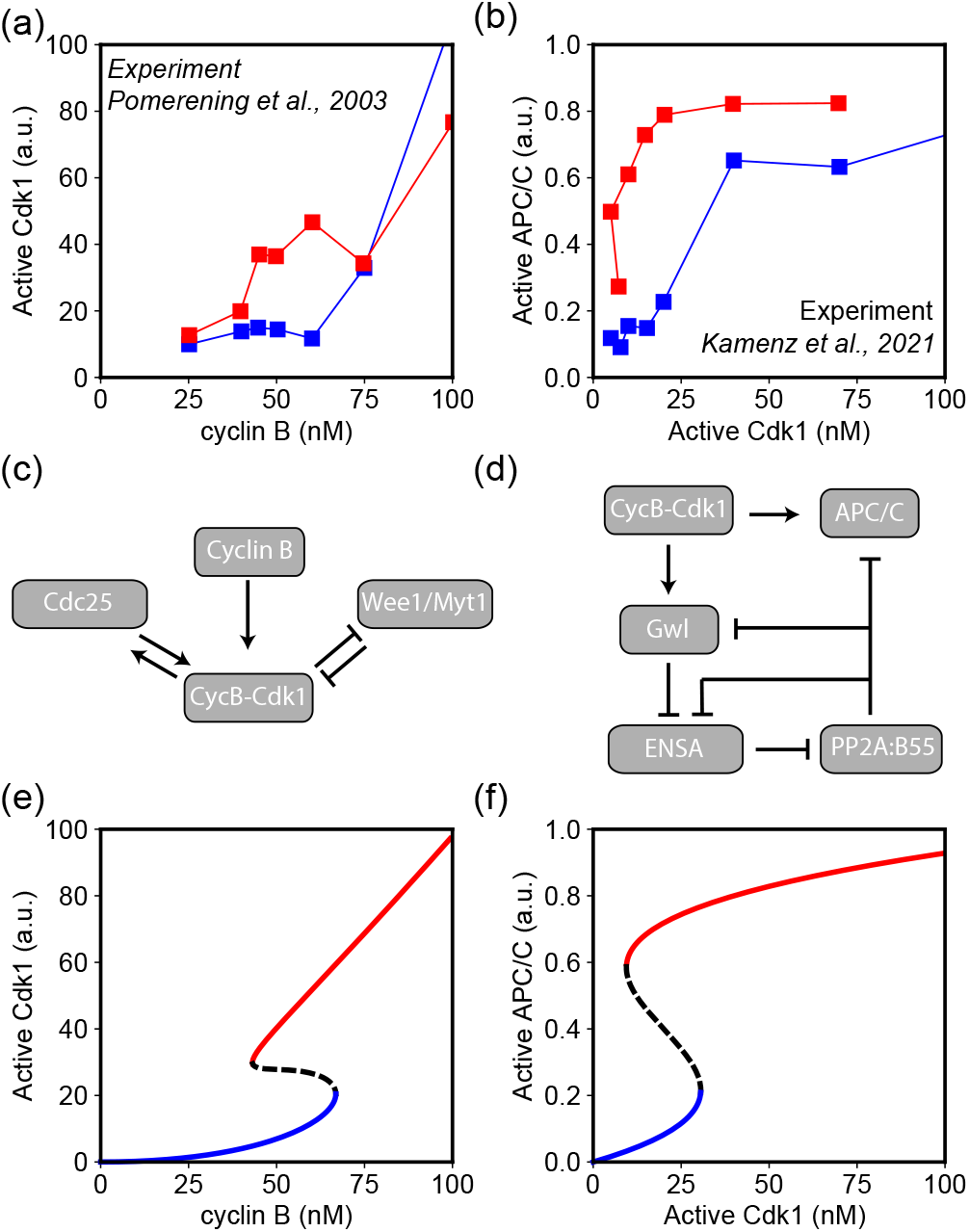
Experimentally measured bistable dose-response curves (a)-(b), their corresponding cell cycle regulatory networks (c)-(d), and their approximations using Eq. (5) in (e)-(f), respectively. (a) Experimentally measured Cdk1 activity (a.u.) in function of cyclin B concentration (nM) Pomerening et al. (2003). (e) Approximated response curve using (*b, C*_*x*_, *C*_*y*_) = (1.88, 55nM, 0.5) as parameters. (b) Experimentally measured APC/C activity (a.u.) in function of active cyclin B - Cdk1 (nM) Kamenz et al. (2021). (f) Approximated response curve using (*b, C*_*x*_, *C*_*y*_) = (1.95, 20nM, 0.5) as parameters.

Here, we theoretically investigate how oscillations emerge by combining negative feedback with one or multiple interlinked bistable switches. We only consider deterministic systems and do not explicitly implement the various molecular interactions that lead to the different bistable switches (those have been studied elsewhere, see e.g. Novak and Tyson (1993); Pomerening et al. (2003); Sha et al. (2003); Mochida et al. (2016); Rata et al. (2018); Kamenz et al. (2021); De Boeck et al. (2021)). Instead, we approximate each experimentally measured bistable response curve by using a single cubic function [see Figure 1(e)-(f)]. We then use them to study how oscillations arise from negative feedback in combination with one or two such bistable switches of varying shapes. In doing so, we show that having two interlinked bistable switches lead to oscillations over a wider range of parameters, thus increasing their robustness.

## Results

### A generic cell cycle oscillator model based on experimentally measured bistable response curves

#### Approximating bistable response curves

We choose a cubic function *x* = *f* (*y*) = *y* −*by*^2^ + *y*^3^ as one of the simplest functions to describe an S-shaped curve. The turning points (folds *F*_1_ and *F*_2_) of this S-shaped response curve can be found by solving d*f* (*y*)*/*d*y* = 0:

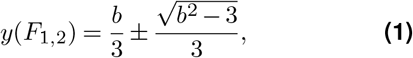

which shows that the system becomes S-shaped when 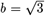, and the center *C* of the S-shaped region is given by:

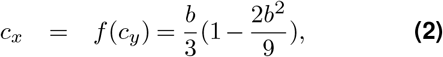

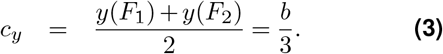

The whole S-shaped region (going from *F*_1_ to *F*_2_) thus scales with *b*. To approximate experimentally measured dose-response curves, we introduce the transformation (*x, y*) = (*c*_*x*_*X/C*_*x*_, *c*_*y*_*Y/C*_*y*_), thus re-positioning the center of the S-shaped curve from (*x, y*) = (*c*_*x*_, *c*_*y*_) to the point (*X, Y*) = (*C*_*x*_, *C*_*y*_). In Figure 2(a), such response curves are plotted for varying values of *b* using *C*_*x*_ = 50 and *C*_*y*_ = 0.5. In this case the left fold *F*_1_ touches the *y*-axis (*x* = 0) when *b* = 2. Figure 2(b) shows the same response curves, but now multiplied by *X*. The reasoning is that, in the context of cell cycle regulation, the red response curves in Figure 2(a) could be used to approximate the fraction (from 0 to 1) of active Cdk1 (or APC/C) in function of cyclin B (or active Cdk1), while the response curve in Figure 2(b) could be interpreted as the total amount of active Cdk1 in function of cyclin B levels. The blue curves are thus the red ones multiplied by the amount of cyclin B, where we use the realistic assumption that all cyclin B proteins quickly bind to Cdk1 to form a complex.

**Figure 2.**
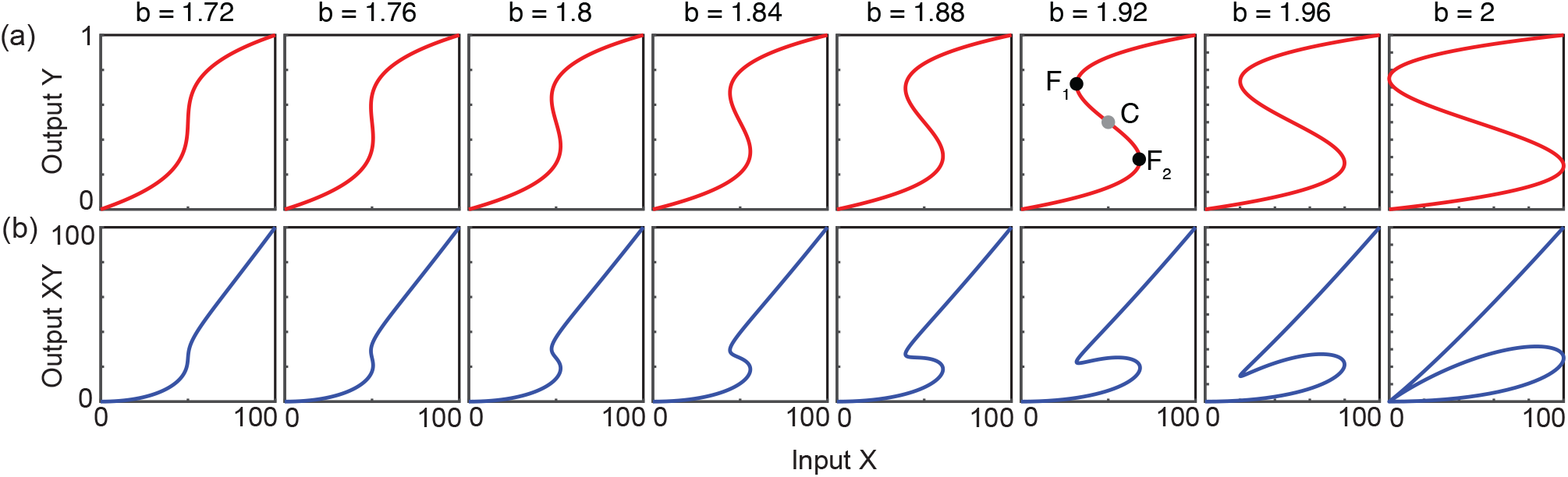
(a) Input (X) - Output (Y) response curves given by Eq. (5) with *C*_*x*_ = 50 and *C*_*y*_ = 0.5 for increasing bistability by increasing the parameter *b*. (b) The same response curves as in (a), but now the output Y is multiplied by the input X.

Figure 1(a)-(b) shows experimentally measured bistable dose-response curves, where the blue markers indicate the measured values when increasing the input, while the red markers depict the measurements when subsequently decreasing the input again. In 2003, Pomerening *et al*. controlled the concentration of (non-degradable) cyclin B in frog egg extracts, and then proceeded to measure the activity of cyclin B - Cdk1 complexes Pomerening et al. (2003) [Figure 1(a)]. This approach allowed the authors to reveal the presence of hysteresis in the system, where for cyclin B concentration between approx. 40 nM and 70 nM the system could be in two possible steady states: low Cdk1 activity (corresponding to interphase) or high Cdk1 activity (corresponding to mitotic phase). Similar experiments were carried out by Sha *et al*. around the same time Sha et al. (2003). We chose (*b, C*_*x*_, *C*_*y*_) = (1.88, 55nM, 0.5) by hand to visually approximate the experimentally measured curve [Figure 1(c)]. While this approximation by the cubic expression is not able to capture the finer details of the measured response curve, it allows to describe the region of bistability. More recently, Kamenz *et al*. used similar techniques in frog egg extracts to measure the activity of APC/C by quantifying how quickly fluorescently-labeled securin was degraded by the proteasome (triggered by APC/C through ubiquitination) in the presence of varying levels of active Cdk1 complexes Kamenz et al. (2021). These experiments once again revealed bistability of APC/C activity in function of Cdk1 activity [Figure 1(b)]. Similarly as before, using (*b, C*_*x*_, *C*_*y*_) = (1.95, 20nM, 0.5), we obtain a good approximation of this response curve [Figure 1(d)].

In order to turn the *S*-shaped dose-response curves in Figure 2 into a dynamical bistable system, we introduce the following ordinary differential equation (ODE) which ensures that the system will approach the steady state solutions given by the cubic function, and a parameter *ϵ* that controls how fast the system will relax to the steadystate solution:

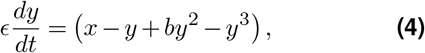

or when re-positioned to the point (*X, Y*) = (*C*_*x*_, *C*_*y*_) as depicted in Figures 1 and 2, the ODE becomes the following:

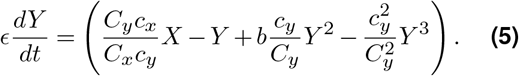

Note that in this work sometimes we use the term “bistable” and “S-shaped” interchangeably for ease of notation. However, we want to emphasize that strictly speaking having an S-shaped dose-response curve does not imply that the overall dynamical system behaves in a bistable way. The dynamical system (4) does exhibit bistability for a range of input values *x*, but this depends on the details of the dynamical system built on such S-shaped response curves. In particular, we will often refer to combining “bistable” response curves to study oscillator properties. In such case, we refer to the fact that a small part of the interaction network can function in a bistable manner in the absence of negative feedback, but the system as a whole behaves as an oscillator when adding negative feedback through APC/Cmediated cyclin B degradation.

#### Turning bistable response curves into a dynamical cell cycle oscillator model

How can we now use such equations that capture bistability in different parts of the cell cycle regulatory network to create a model of cell cycle oscillations? First, we introduce an ODE to describe the synthesis and degradation of cyclin B (*X*), see Eq. (6). Cyclin B is synthesized at a constant rate (*k*_*s*_ [nM/min]), where this synthesis rate is experimentally known to be around 1 nM/min (see Table 1 and Refs. Chang and Ferrell Jr (2013); Yang and Ferrell (2013)). Furthermore, it is degraded at a fixed basal level (*d*_1_ [min^−1^]), as well as in a APC/C (Z) - dependent manner (*d*_2_ [min^−1^]). Here, experiments have motivated the APC/C-driven degradation rate *d*_2_ to be approx. in the range 0.05 - 0.5 min^ −1 Rombouts et al. (2018); Yang and Ferrell (2013), and *d*_1_ should be much smaller (exact value of *d*_1_ not critical for the dynamics discussed in this work). Second, in Eq. (7), we capture the experimental observation of bistability in Cdk1 activity in response to fixed values of cyclin B concentration, see Figure 1(a),(c),(e). *X* corresponds to cyclin B concentration, and *Y* is the fraction of active Cdk1 complexes. For fixed values of cyclin B concentration (*X*), the system relaxes to the steady state solution shown in Figure 1(e). Third, in Eq. (8), APC/C activity (*Z*) is controlled by Cdk1 activity (*X ×Y*). So, the variables are defined as follows:

**Table 1.** Parameters used in the different cell cycle oscillator models: built on cyclin B (X) - Cdk1 (XY) bistability (13)-(14), built on Cdk1(XY) - APC/C (Z) bistability (15)-(16), and built on the two interlinked bistable switches (6)-(8).

| Par. | Meaning | (13)-(14) | (15)-(16) | (6)-(8) | References |
| --- | --- | --- | --- | --- | --- |
| $k_s$ | cyclin synthesis rate | 1 nM/min | 2.3 nM/min | 1 nM/min | Chang and Ferrell Jr (2013) |
| $d_1$ | basal degradation rate | 0.001 min<sup>-1</sup> | 0.001 min<sup>-1</sup> | 0.001 min<sup>-1</sup> | Chang and Ferrell Jr (2013) |
| $d_2$ | APC/C-driven degradation rate | 0.1 min<sup>-1</sup> | 0.1 min<sup>-1</sup> | 0.05 min<sup>-1</sup> | Rombouts et al. (2018) |
| $b_y$ | controls bistability range | 1.88 | - | 1.88 | Pomerening et al. (2003) |
| $b_z$ | controls bistability range | - | 1.9 | 1.95 | Kamenz et al. (2021) |
| $\epsilon_y$ | relaxation time | 0.02 min | - | 0.02 min | This work |
| $\epsilon_z$ | relaxation time | - | 0.02 min | 0.1 min | This work |
| $n$ | Hill exponent | 20 | - | - | Yang and Ferrell (2013) |
| $C_x$ | location switch in X | 55 nM | - | 55 nM | Pomerening et al. (2003) |
| $C_y$ | location switch in Y | 0.50 | - | 0.50 | Pomerening et al. (2003) |
| $C_{xy}$ | location switch in XY | 27 nM | 50 nM | 20 nM | Kamenz et al. (2021) |
| $C_z$ | location switch in Z | - | 0.50 | 0.50 | Kamenz et al. (2021) |

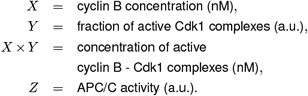

Their dynamical evolution is controlled by this set of ODEs:

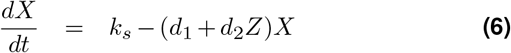

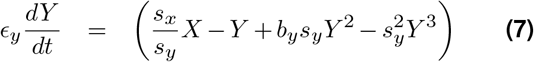

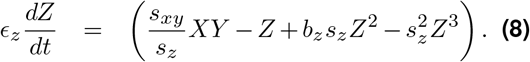

using the following scaling factors *s*_*x*_, *s*_*y*_, *s*_*z*_, *s*_*xy*_:

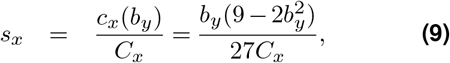

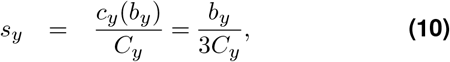

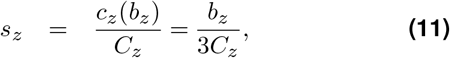

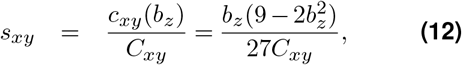

Using the system parameters given in Table 1, we find oscillations in (*X, Y, Z*) driving the cell cycle forward. In Figure 3, we modulate the parameter *b*_*y*_ in time from 1.94 to 1.73 to 1.91, mimicking the experimental observation that the *S*-shaped response curve of Cdk1 activity in function of cyclin B concentration is abolished after the first embryonic cell cycle Tsai et al. (2014). As a result, the first cell cycle is long (approx. 75 min), while the next cell cycles are short (approx. 30 min). Finally, after approx. 10 cell cycles, embryonic divisions gradually slow down again, which has been attributed at least partially to increases in *b*_*y*_ (the ratio of enzymatic activity of Wee1 vs. Cdc25 increases). The experimental data of embryonic cell division timing at 21-23°C plotted (in red) in Figure 3(c) is taken from Satoh (1977) (with the data of the first cell cycle our own unpublished data). Figure 3 thus shows that tuning the width of the underlying bistable response curves allows to change the experimentally observed changes in period of the cell cycle oscillator. While most parameters in Table 1 are experimentally well motivated, the timescale parameters *ϵ*_*y*_ and *ϵ*_*z*_ are less well known and getting the results in Figure 3 required *ϵ*_*y*_ and *ϵ*_*z*_ to be sufficiently small. *ϵ*_*y*_ is mainly determined by the phosphorylation and dephosphorylation rates of Cdk1 by Wee1 and Cdc25 (and vice versa). The absolute value of these rates are experimentally not well known (see discussion in Ref. Yang and Ferrell (2013)). The other timescale determining the activation of APC/C by Cdk1 is given by *ϵ*_*z*_, and has been measured to be time-delayed (with a delay time of minutes) Yang and Ferrell (2013); Rombouts et al. (2018). To choose values for (*ϵ*_*y*_, *ϵ*_*z*_), we carried out simulations over a wide range of parameters, analyzing the resulting cell cycle oscillation period for each parameter set. In doing so, we found that *ϵ*_*y*_ needs to be smaller than approx. 10−2, while *ϵ*_*z*_ needs to be smaller than approx. 10−1 to obtain realistic oscillation periods. Having sufficiently small values of *ϵ*_*y*_ and *ϵ*_*z*_ (see Table 1) also allows for a clear interpretation of the dynamics as relaxation-like oscillations.

**Figure 3.**
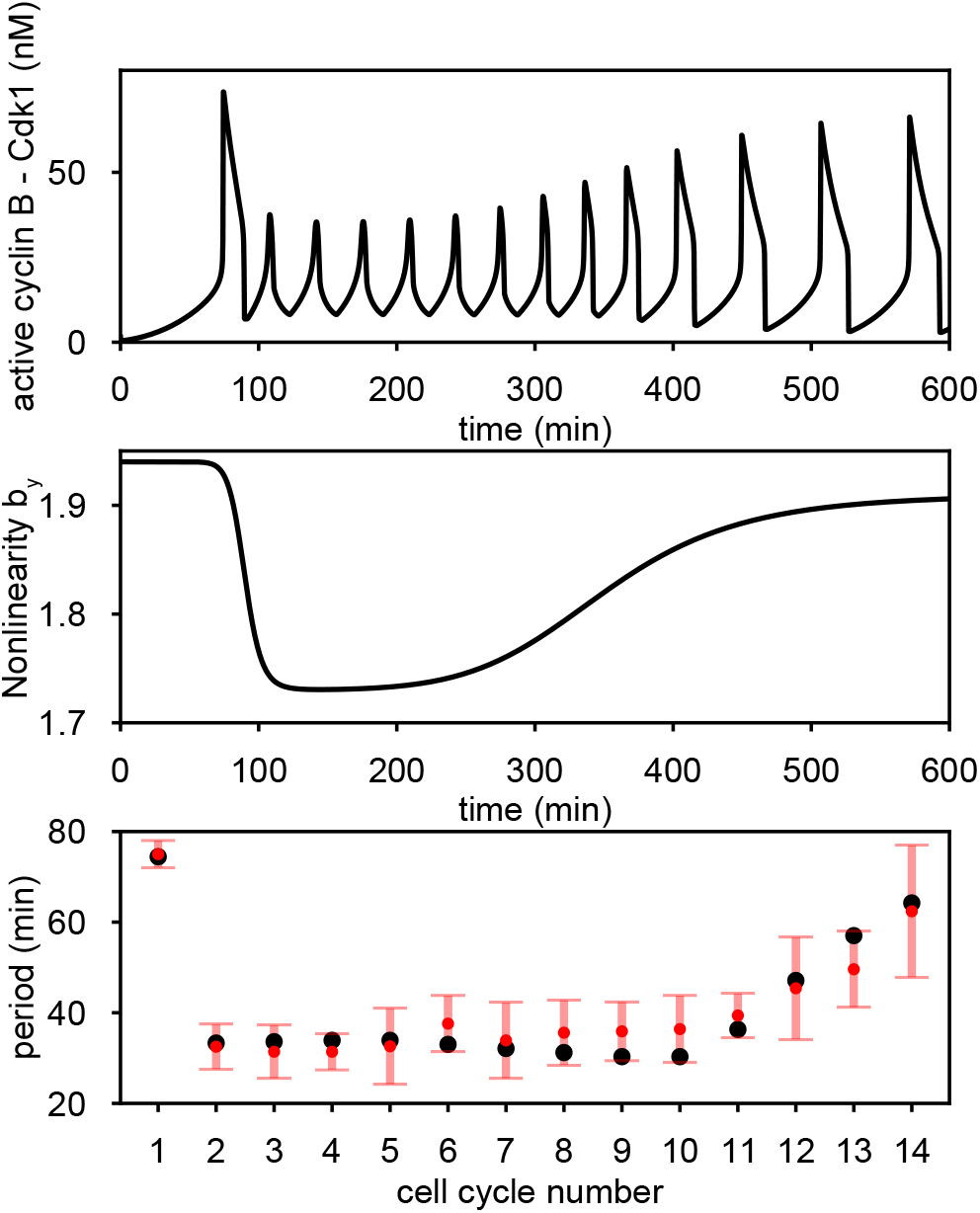
(a) Simulation of the model for the cell cycle oscillator built on two interlinked bistable switches. (b) In time *b*_*y*_ was changed from 1.94 to 1.73 to 1.91. (c) The detected period for every consecutive cell cycle: in black for the simulated time series in (a), and in red for the experimentally measured periods (mean period *±* one standard deviation) Satoh (1977); Anderson et al. (2017)

In what follows, we will introduce and elaborately study the dynamics of different oscillator models built on each of the bistable switches in Figure 1, as well as both switches combined. Note that we introduced a similar approach in De Boeck et al. (2021), which allowed for greater control over the response curve. However, here we limit ourselves to a cubic nonlinear function as it is likely one of the simplest expressions that can be used to reproduce a bistable switch.

#### A cell cycle oscillator built on cyclin B - Cdk1 bistability

A cell cycle oscillator model can be constructed by taking into account the experimental observation of bistability in Cdk1 activity in response to fixed values of cyclin B concentration, see Figure 1(a),(c),(e). To do so, we study a simplified version of Eq. (6)-(8). We complement Eq. (7) with a modified version of Eq. (6), assuming that APC/C activity (*Z*) is a nonlinear, ultrasensitive function of Cdk1 activity as measured in Yang and Ferrell (2013); Tsai et al. (2014):

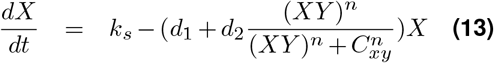

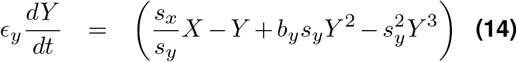

The parameters are given in Table 1, where additionally the scaling factors *s*_*x*_ and *s*_*y*_ are given by (9)-(12). Using the parameters from Table 1, Figure 4(a) shows the time evolution of cyclin B concentration (*X*) in blue and the corresponding concentration of active Cdk1 (*X ×Y*) in orange. This simple model thus reproduces cell cycle oscillations with a realistic period of around 30 minutes, where Cdk1 is periodically sharply activated and inactivated.

**Figure 4.**
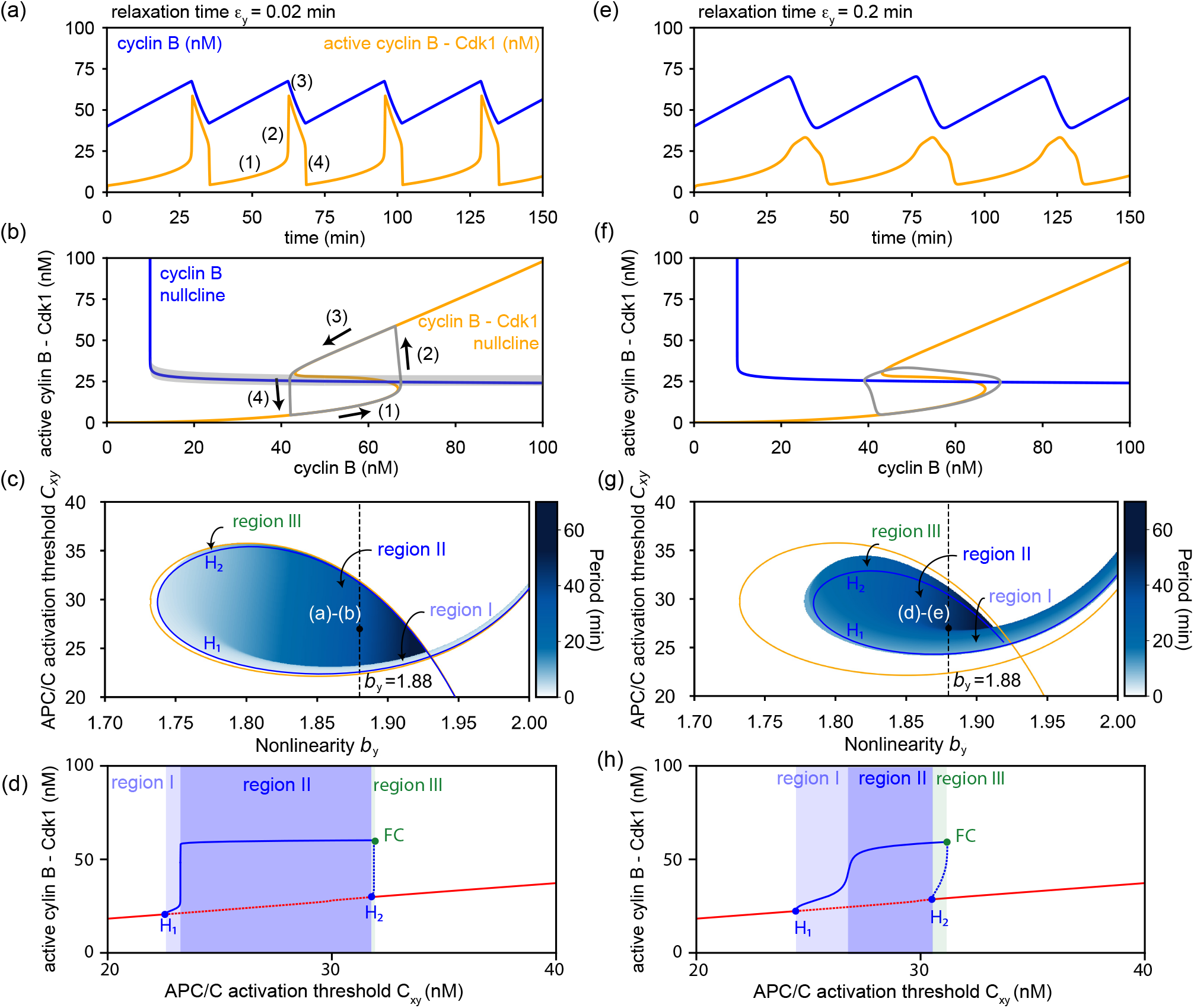
Simulation of the model for the cell cycle oscillator built on cyclin B - Cdk1 bistability (13)-(14). Time series (a,e) and phase space (b,f). Phase diagram in *b*_*y*_ and *C*_*xy*_ (c,g) showing the period of the oscillation in blue color map, the Hopf bifurcations (blue line) and the intersection of the cyclin B nullcline with the folds of the cyclin B - Cdk1 nullcline (orange line). Bifurcation diagram in *C*_*xy*_ (d,h) for *b*_*y*_ = 1.88 showing the value of active cyclin B - Cdk1 of the stationary state (in red) and the maximum of the oscillation (in blue). The Hopf bifurcations are indicated by H_1,2_ and the Fold of cycles by FC.

The functioning of this oscillator can be visually interpreted in the phase space (X, X *×*Y) = (cyclin B, active Cdk1) by looking at the nullclines (curves where *dX/dt*=0 or *dY/dt*=0) in 4(b). The Cdk1 (*Y*) - nullcline (*dY/dt*=0) in orange shows the typical *S*-shaped curve, and the cyclin B (*X*) - nullcline (*dX/dt*=0) in blue illustrates the ultrasensitive activation of APC/C when Cdk1 activity crossed the threshold *C*_*xy*_ = 27 nM. The cell cycle oscillations plotted in Figure 4(a) correspond to a limit cycle attractor in this phase space, plotted in gray in Figure 4(b). The oscillations consist of four phases [indicated as (1)-(4)]:

1. Cyclin B is synthesized at a rate *k*_*s*_, while APC/C is inactive, and the system moves along the bottom branch of the *S*-shaped curve;
2. The system reaches the turning point of the *S*-shaped curve, at which point Cdk1 is quickly (if *ϵ*_*y*_ is sufficiently small) activated and the system jumps to the top branch of the *S*-shaped curve;
3. Active Cdk1 activates APC/C, cyclin B is degraded at a rate dominated by *d*_2_, such that the system moves down along the top branch of the *S*-shaped curve;
4. Finally, the left turning point of the *S*-shaped curve is reached and the system jumps down to the bottom branch again, thus inactiving APC/C such that the system is reset and phase (1) starts over.

Notice that the blue cyclin B - nullcline intersects the middle branch of the orange *S*-shaped Cdk1 - nullcline. For small values of *ϵ*_*y*_, this serves as a condition for obtaining relaxation-type oscillations in the system. This visual interpretation allows us to easily see when cell cycle oscillations exist in this model. For example, by changing the APC/C activation threshold *C*_*xy*_ the cyclin B - nullcline can be shifted up or down. When *C*_*xy*_ is changed outside of the approximate interval [22 nM, 32 nM] [the gray region in Figure 4(b)], it will intersect the top or bottom branch of the *S*-shaped nullcline, leading to a stable steady-state solution. This can also be seen in Figure 4(c) where we show the region of oscillations (and their period) in the parameter space (*b*_*y*_, *C*_*xy*_). Here, the orange lines indicate the parameter values for which the cyclin B - nullcline intersects either turning point of the *S*-shaped Cdk1 - nullcline. As expected, cell cycle oscillations that encircle the *S*-shaped Cdk1 - nullcline have a higher amplitude and period, and they are contained between these orange lines in the parameter space.

When increasing the width of the *S*-shaped Cdk1- nullcline further (by increasing *b*_*y*_), the oscillation period increases and the parameter region of oscillations decreases. This makes sense as it becomes increasingly difficult for the cyclin B - nullcline to intersect the middle branch of the *S*-shaped nullcline. Relaxation oscillations of higher amplitude and period disappear entirely above *b*_*y*_ ≈1.93, when both turning points of the *S*-shaped nullcline have the same y-value: *X*(*F*_1_) *Y ×* (*F*_1_) = *X*(*F*_2_) *×Y* (*F*_2_). Note that there still exists a narrow region of fast, small amplitude oscillations above this critical point. These oscillations correspond to a limit cycle that does not encircle the top and bottom branch of the *S*-shaped nullcline.

From a dynamical systems perspective, the cell cycle oscillations originate from a Hopf bifurcation. When constructing a bifurcation diagram for changing values of the APC/C activation threshold (*C*_*xy*_), we find that the steady state solutions (in red) lose their stability at two Hopf bifurcations H_1_ and H_2_ [see Figure 4(c)-(d)]. The maxima of the resulting oscillations are plotted in blue. Considering Cdk1 activity is low when the steady state solution lies on the lower branch of the bistable switch and Cdk1 activity is high when on the upper branch, the low activity steady state loses its stability in a supercritical bifurcation (H_1_), while the high activity steady state solution stabilizes in a subcritical bifurcation (H_2_) when increasing the APC/C activation threshold (*C*_*xy*_). Beyond H_1_, the oscillations are initially of low amplitude and period (region I), but then sharply increase in amplitude and period when increasing *C*_*xy*_ (region II). Such sharp transitions are referred to as Canard behavior, or Canard-explosion, typical in fast-slow systems Desroches et al. (2012); Kuehn (2015). As H_2_ is a subcritical bifurcation, there exists a (narrow) region in parameter space (between H_2_ and a fold of cycles FC, region III) where stable oscillations coexist with a stable high activity steady state solution.

How do the cell cycle oscillations change when keeping the nullclines fixed, but changing the relaxation time *ϵ*_*y*_ and the rates of cyclin synthesis *k*_*s*_ and APC/Cdriven degradation *d*_2_? The visual interpretation using the location of intersecting nullclines as a way to predict the presence of cell cycle oscillations requires sufficient timescale separation. When *ϵ*_*y*_ increases (i.e., the system is no more on the singular perturbation regime) this is no longer valid. In this limit, the Canard explosion can be defined in terms limit cycle’s curvature change, and detected using the inflection-line method Brøns and Bar-Eli (1997); Desroches and Jeffrey (2011).

In Figure 4(e)-(h) we repeat the same analysis as in Figure 4(a)-(d), but now for *ϵ*_*y*_ = 0.2 min, where the oscillations are less relaxation-like and thus more sinusoidal [Figure 4(e)-(f)]. Here, oscillations overall exist in a smaller region in parameter space, but they do persist beyond *b*_*y*_≈ 1.93, which was a critical point when the time scale separation was sufficiently large [Figure 4(g)]. When increasing *ϵ*_*y*_ even further, oscillations are lost for these parameters, even though the cyclin B - nullcline still intersects the middle branch.

In Figure 5, we illustrate how the rates of cyclin synthesis *k*_*s*_ and APC/C-driven degradation *d*_2_ affect the cell cycle oscillations. As the synthesis and degradation rates increase, the period of the oscillations decreases [Figure 5(a)]. Furthermore, oscillations only exist within a well-defined region of synthesis and degradation rates. For a realistic value of the degradation rate (*d*_2_ = 0.1 min^−1^), Figure 5(b) illustrates in more detail how the oscillations change with the cyclin synthesis rate *k*_*s*_. At high *k*_*s*_, oscillations originate in a Hopf bifurcation (H_1_) that shows a fast Canard increase in period and amplitude (see inset). For lower values of *k*_*s*_ the oscillations disappear in a homoclinic bifurcation (indicated by the black dots). The oscillation period increases (in principle towards ∞) when approaching the homoclinic bifurcation. For larger values of the degradation rate *d*_2_, the oscillations become more sinusoidal, faster and of lower amplitude. Here, for low synthesis rates *k*_*s*_, stable oscillations are born in a supercritical Hopf bifurcation (H_2_) [see Fig. 5(c)]. The oscillations are then lost at higher values of the synthesis rate *k*_*s*_ in either a fold of cycles (FC) where the stable limit cycle touches the unstable limit cycle born in a subcritical Hopf bifurcation H_3_; or in another supercritical Hopf bifurcation (H_3_). Note that the Canard cycles that originate at the Canard explosion shown in Fig. 5(c) for *d*_2_ = 1 min^−1^ may persist when decreasing *d*_2_ [see Fig. 5(c) for *d*_2_ = 0.01 min^−1^], leading to a homoclinic connection with a Canard segment. The Hopf and saddle-node bifurcations, as well as the fold of cycles, are plotted in Figure 5(a), where the oscillations are bounded by the supercritical Hopf bifurcations, the fold of cycles and/or the homoclinic bifurcations (the homoclinic bifurcation line is not shown). Overall, the dynamics unfold from two Takens-Bogdanov (TB) points of co-dimension two (in purple). Here, the saddle-node bifurcation, Hopf bifurcation, and homoclinic bifurcations come together, while the fold of cycles (FC) also occurs very close to it.

**Figure 5.**
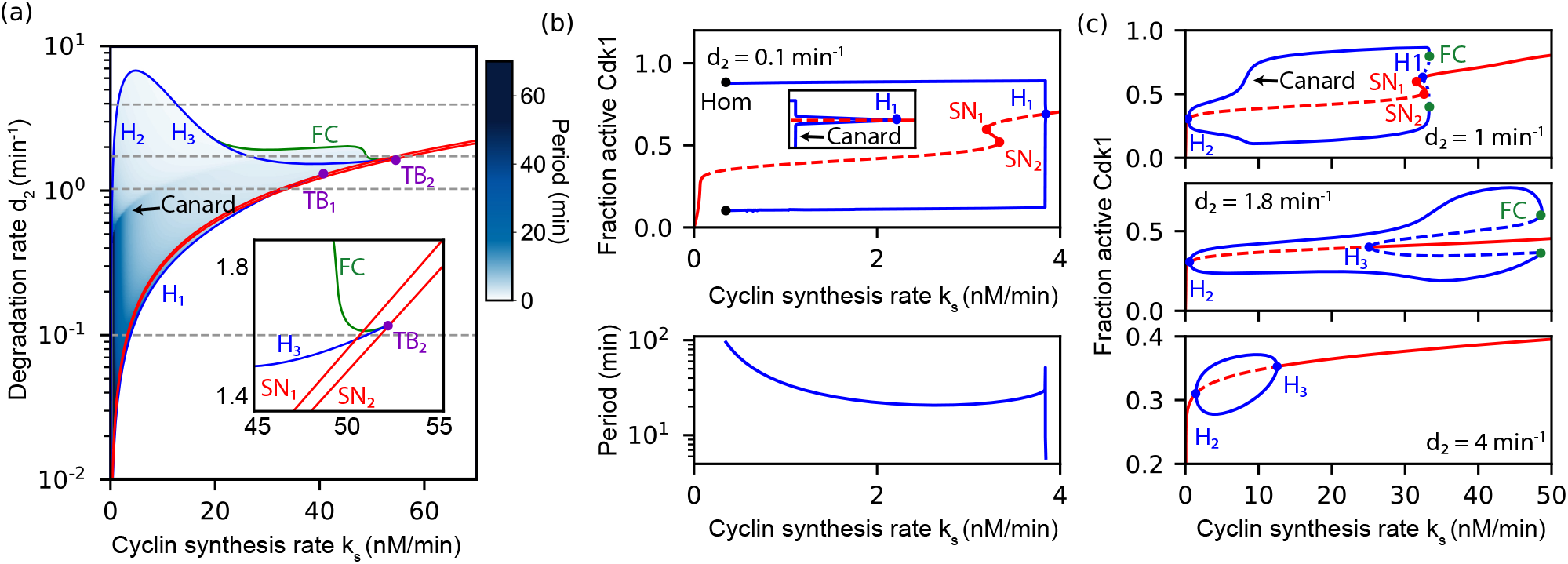
(a) Phase diagram of model (13)-(14) in the parameter space (*k*_*s*_, *d*_2_). Different bifurcations are indicated: Hopf bifurcations (H_1_, H_2_, H_3_ - blue), Folds of cycles (FC - green), and saddle-node bifurcations (SN_1_, SN_2_ - red). TB_1_ -TB_2_ are two co-dimension two Takens-Bogdanov points. When stable oscillations exist, their period is also indicated in a blue colorscale. (b)-(c) Bifurcation diagrams in function of cyclin synthesis *k*_*s*_ for different values of the degradation rate *d*_2_. All other parameters are given in Table 1.

#### A cell cycle oscillator built on Cdk1 - APC/C bistability

A recent series of experiments Mochida et al. (2016); Rata et al. (2018); Kamenz et al. (2021) demonstrated that there also exist bistable response functions elsewhere in the cell cycle regulatory network, i.e. in the central regulation of APC/C activity by active Cdk1 [see Figure 1(b)]. Starting from Eqs. (6)-(8), we omit Eq. (7) by assuming that cyclin B quickly binds to Cdk1, and that each such cyclin B - Cdk1 complex is in its active state (*Y* = 1) and *X ×Y* = *X*. We then choose a parameter set (see Table 1) to approximate this experimentally measured response curve of APC/C activity (*Z*) in function of the concentration of active Cdk1 (*X ×Y*), see Figure 1(d). This leaves us with the following dynamical cell cycle control system:

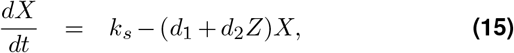

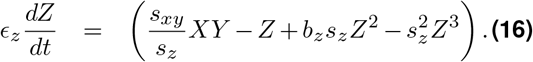

The control parameters are given in Table 1. Note that the parameters to describe the bistable response curve are different than the ones found to fit the experimentally measured curve in Figure 1(d). This was required to obtain oscillations in this model and will be discussed later in this section. In order to obtain oscillations in this model the parameters need to be different to compensate for the absence of the (cyclin B, active Cdk1) switch as can be observed in phase diagrams shown in Figures 6(c) and 7(a).

**Figure 6.**
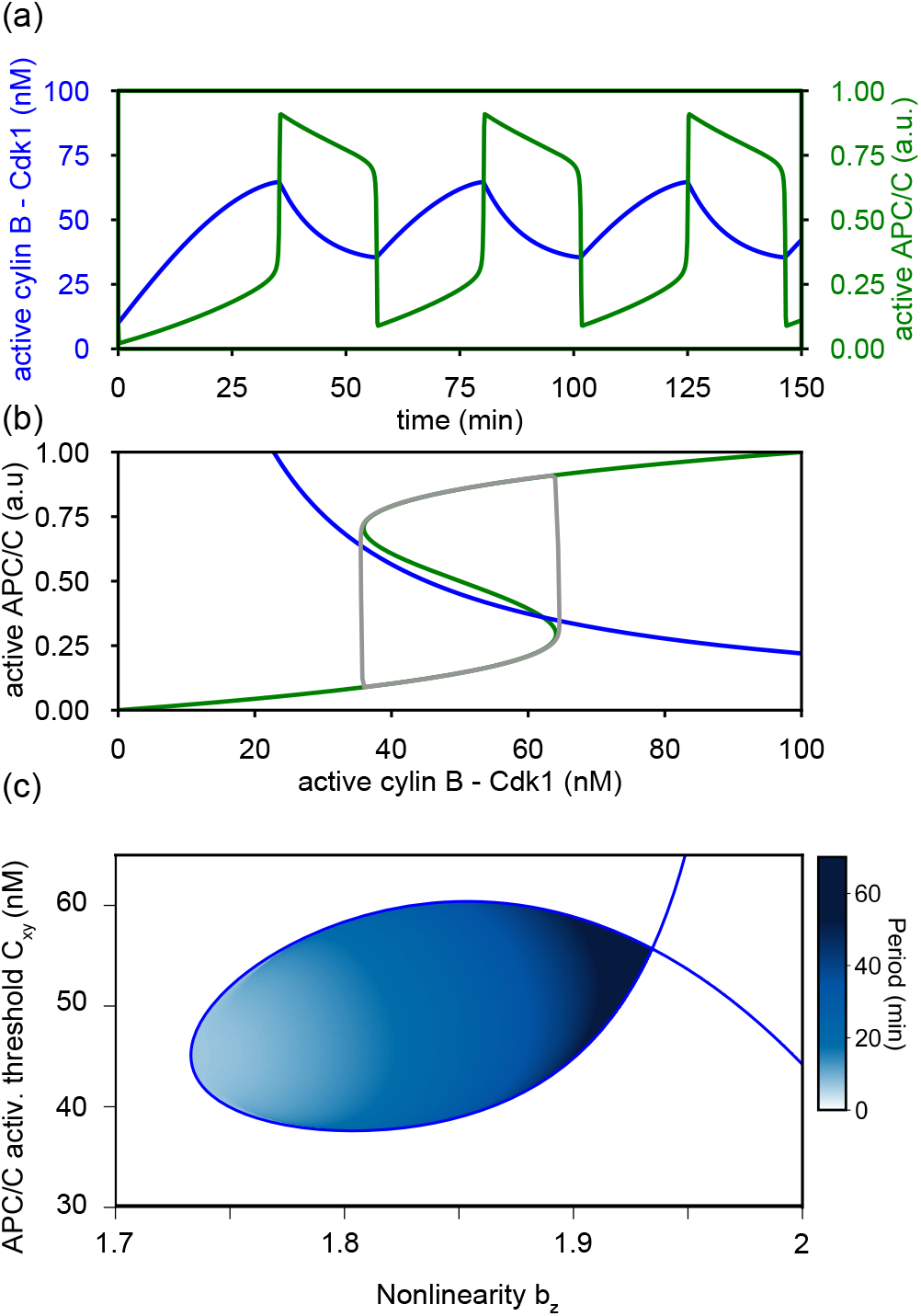
(a) Simulation of model (15)-(16) for the cell cycle oscillator built on Cdk1 - APC/C bistability: time series and phase space. (b) (*b*_*z*_, *C*_*xy*_)-phase diagram with Hopf bifurcations in blue. Parameters are given in Table 1

**Figure 7.**
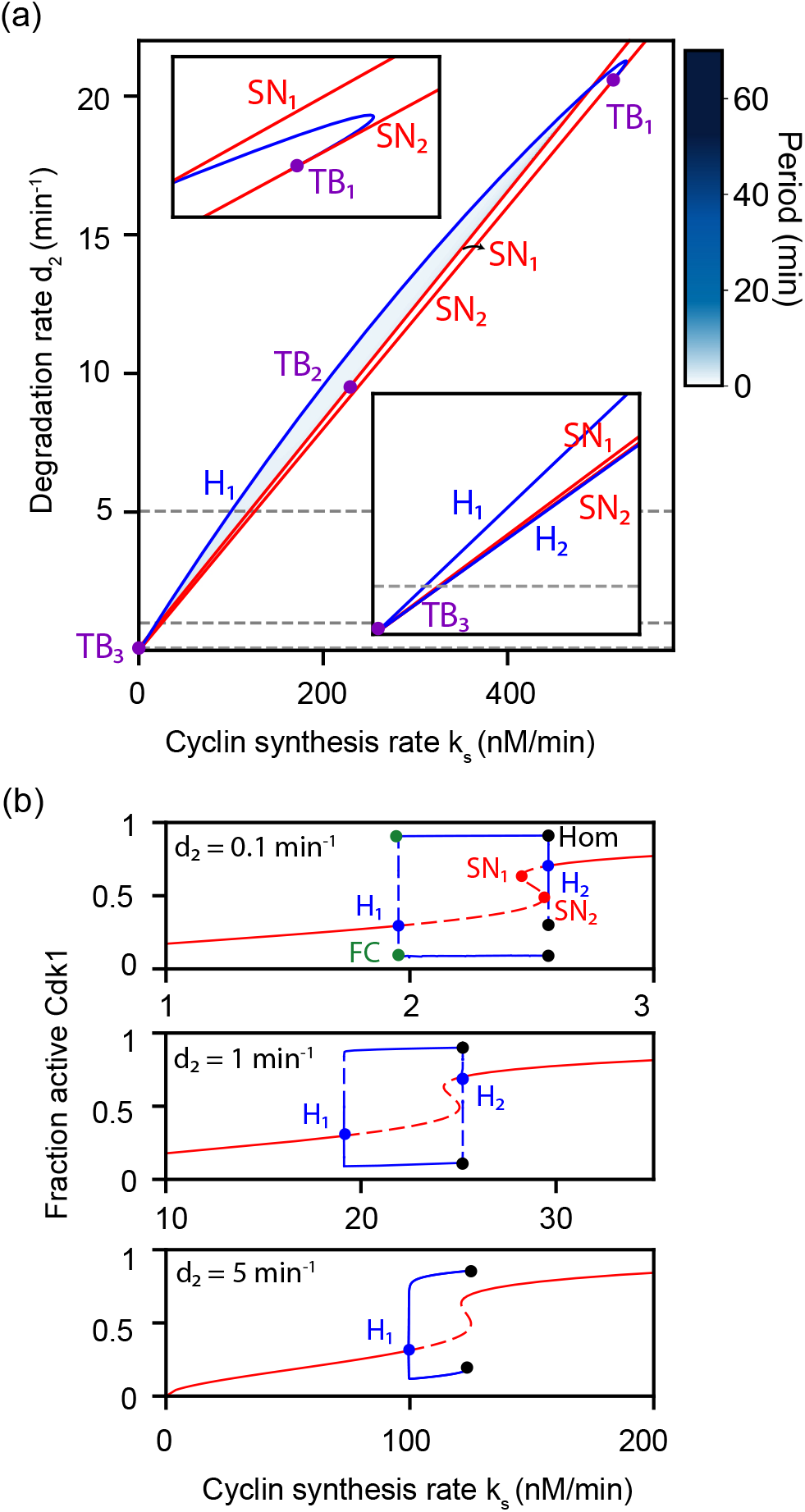
(a) Phase diagram of model (15)-(16) in the parameter space (*k*_*s*_, *d*_2_). Different bifurcations are indicated: Hopf bifurcations (H_1_ - blue), Fold of Cycles (FC - green), and saddle-node bifurcations (SN_1_, SN_2_ - red). TB_1_ - TB_2_ -TB_3_ are three co-dimension two Takens-Bogdanov points. When stable oscillations exist, their period is also indicated in a blue colorscale. (b) Bifurca- tion diagrams in function of cyclin synthesis *k*_*s*_ for *d*_2_ = 0.1, 1, and 5 min^−1^. All other parameters are given in Table 1.

Oscillations in the activity of Cdk1 and APC/C are shown in Figure 6. While the oscillations look very similar to the ones in Figure 4(a)-(b), there are important differences. The oscillations are also relaxation-like, but here it is not Cdk1 (blue) which is periodically sharply activated and deactivated, but rather the activity of APC/C (green). Indeed, it is now the APC/C-nullcline (*dZ/dt* = 0) that is *S*-shaped in the phasespace (*X× Y, Z*) = (active Cdk1, active APC/C), in contrast with model (13)-(14) where active Cdk1 was an *S*-shaped function of cyclin B. Another difference is that the cyclin nullcline (*dX/dt* = 0) is no longer ultrasensitive [as in model (13)-(14)]. Despite these differences, the origin of oscillations can be similarly interpreted in the phase space in Figure 4(b). Oscillations exist when the cyclin nullcline (blue) intersects the middle branch of the *S*-shaped APC/C nullcline (green), provided that the relaxation time *ϵ*_*z*_ is sufficiently small ensuring that APC/C is quickly activated/deactivated.

Similar as in the previous section, the oscillations originate from a Hopf bifurcation, shown by the blue line in Figure 4(b), illustrating how the nonlinearity *b*_*z*_ (controlling the width of the *S*-shaped APC/C response curve) and *C*_*xy*_ (controlling the threshold of APC/C activation) affect the region of oscillations. Although the *S*-shaped response curve relates to different variables in models (13)-(14) and (15)-(16), both show similar dynamical properties. Figure 7(a) shows the region of oscillations when changing the cyclin synthesis rate *k*_*s*_ and degradation rate *d*_2_. Representative bifurcation diagrams for different values of *d*_2_ are plotted in Figure 7(b). An unstable limit cycle is created in a subcritical Hopf bifurcation (H_1_), which is stabilized in a fold of cycles (FC). These limit cycle oscillations disappear at larger *k*_*s*_ in a homoclinic bifurcation (Hom). Several such homoclinic bifurcations, as well as a fold of cycles and another Hopf bifurcation H_2_, occur close to the saddle-node bifurcation SN_2_. A more detailed bifurcation analysis of this complicated scenario is left for future work. The main Hopf bifurcation and saddle-node bifurcation lines are shown in 7(a) in the (*k*_*s*_, *d*_2_) parameter plane, illustrating that the dynamics again unfold from several co-dimension two TB points.

#### A cell cycle oscillator built on two interlinked bistable switches

Next, we wondered what would happen if we consider the full model (6)- 8) that incorporates both *S*-shaped responses present in models (13)- 14) and (15)- 16) that we studied separately in the two previous sections. Using the parameters given in Table 1, which are based on experimentally measured *S*-shaped response curves (Figure 1), we simulated the dynamics of (6)- 8). Figure 8(a) shows the resulting oscillations in cyclin B concentration (blue), concentration of active Cdk1 (orange), and APC/C activity (green). A first observation is that the oscillations are again relaxation-like as not only cyclin B - Cdk1 is periodically sharply (de)activated (with a relaxation time *ϵ*_*y*_), but also APC/C activity has a similar sharp transition (with a relaxation time *ϵ*_*z*_). The cell cycle model (6)- 8) consists of three ODEs. Therefore, it is no longer possible to interpret all of its dynamics in a 2D phase plane. It is, however, still informative to project the nullclines and the 3D system dynamics in the phase plane (*X, X ×Y*) = (cyclin B, active Cdk1) and in the plane (*X ×Y, Z*) = (active Cdk1, active APC/C). This can be seen in Figure 8(b)-(c), respectively, which reveal both bistable switches that were measured experimentally (Figure 1) as nullclines: the (cyclin B, active Cdk1) switch (in orange - the Cdk1 nullcline: d*Y* /d*t* = 0) and the (active Cdk1, active APC/C) switch (in green - the APC/C nullcline: d*Z*/d*t* = 0). The third nullcline (in blue - the cyclin B nullcline: d*X*/d*t* = 0) requires using either the expression of the APC/C nullcline or the Cdk1 nullcline as well to be able to plot it in the chosen 2D phase planes, which is why we label them as accordingly. The gray lines indicate the projected limit cycle trajectory in the respective phase planes. This reveals that the oscillations are mainly driven by the (cyclin B, active Cdk1) switch (in orange) as the limit cycle closely circles around this switch. This results in oscillations of large amplitude in Cdk1 activity, as Cdk1 is activated [to a value of Cdk1(F_2_(Cdk1))] when crossing the fold of the Cdk1 nullcline [F_2_(Cdk1)]. As this Cdk1 activity is much larger than the right fold of the APC/C nullcline [F_2_(APC)], APC/C is strongly activated, leading to cyclin B degradation and deactivation of both Cdk1 and APC/C, thus resetting the cell cycle oscillator. We can again observe that the different projected nullclines cut each other in the middle branch of each switch. Whereas in our previous two oscillator model this was found to be a good indicator for oscillations, we remark that this should not necessarily be the case here as the system is no longer two-dimensional.

**Figure 8.**
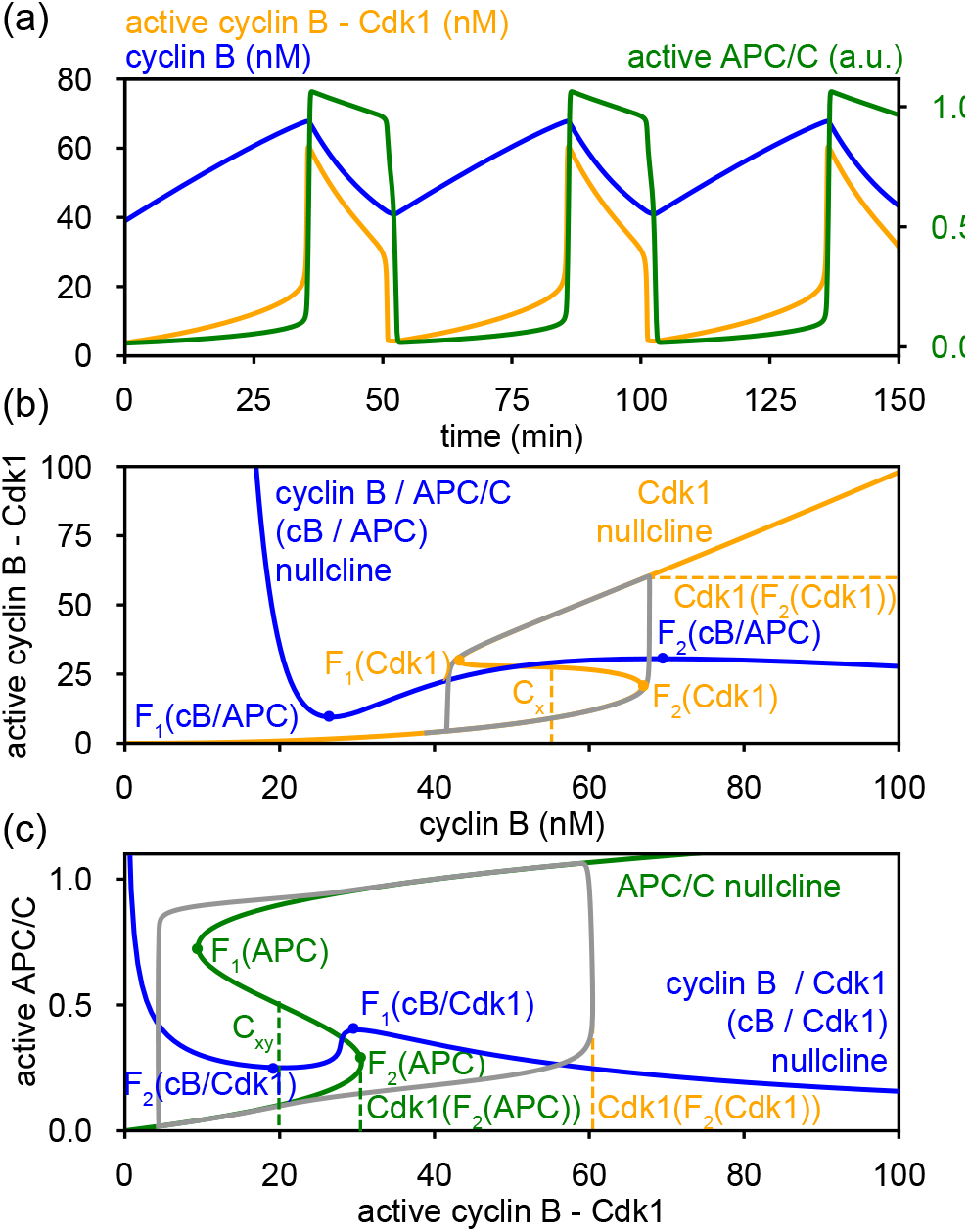
Simulation of the model for the cell cycle oscillator built on two interlinked bistable switches: time series (a) and phase space projections (b,c). Parameters as in Table 1.

We then again explored the influence of the rates of synthesis (*k*_*s*_) and degradation of cyclin B (*d*_2_), see Figure 9. The presence of two bistable switches makes the bifurcation diagrams more complex, now introducing additional saddle-node and Hopf bifurcations. Despite the increased complexity, overall the response when increasing the synthesis rate is similar. Let us more closely consider our standard case from Table 1 where *d*_2_ = 0.05 min^−1^), see last panel in Figure 9(b) and the phase space projections for selected values of *k*_*s*_ in Figure 9(c). For low synthesis rates (e.g. *k*_*s*_ = 0.3 nM/min), the system finds itself in a state of low Cdk1 and APC/C activity. When increasing *k*_*s*_ to about 0.45, the system starts to oscillate at a Hopf bifurcation H_1_. The onset of oscillations is found to occur very close to the saddle-node bifurcation SN_1_, which is the moment when the cyclin B / APC - nullcline crosses the right fold of the Cdk1 nullcline [F_2_(Cdk1)]. Further increasing the synthesis rate then leads to a loss of the oscillations close to the saddle-node bifurcation SN_3_, which occurs when the cyclin B / Cdk1 - nullcline crosses the left fold of the APC/C nullcline [F_1_(APC)]. Beyond SN_3_ two possible states of stationary activity coexist. The dominant state corresponds to high cyclin B - Cdk1 and APC/C activities, while the stationary state with the smallest basis of attraction corresponds to low cyclin B - Cdk1 and high APC/C activities. Increasing *k*_*s*_ even more, only one stable state persists, corresponding to high cyclin B - Cdk1 and APC/C activities. For lower values of the degradation rate *d*_2_ [e.g. *d*_2_ = 0.05, 0.1, 0.5, 1 min^−1^ in Figure 9(b)], the loss of oscillations is found to occur in a homoclinic bifurcation indicated by the black dot close to SN_3_. Another interesting observation is that oscillations exist over an increasingly wide range of synthesis rates *k*_*s*_ as the degradation rate *d*_2_ increases. This contrasts with the disappearance of oscillations for higher degradation rates for the oscillators only built on a single bistable switch, see Figures 5(a) and 7(a).

**Figure 9.**
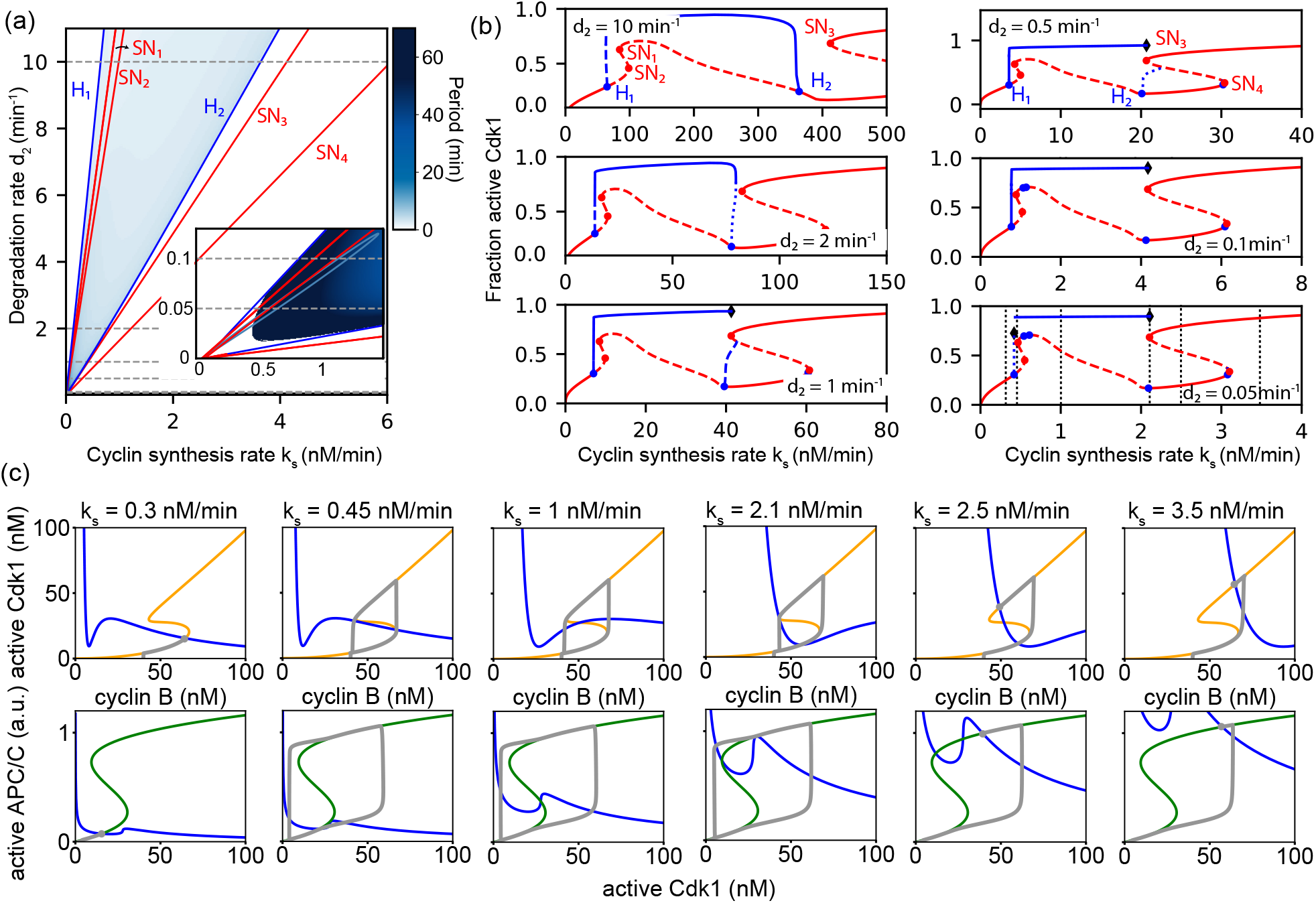
(a) Phase diagram of model (6)-(8) in the parameter space (*k*_*s*_, *d*_2_). Different bifurcations are indicated: Hopf bifurcations (H - blue) and saddle-node bifurcations (SN - red). TB is a co-dimension two Takens-Bogdanov point. When stable oscillations exist, their period is also indicated in a blue colorscale. (b) Bifurcation diagrams in function of cyclin synthesis *k*_*s*_ for *d*_2_ = 0.05, 0.1, 0.5, 1, 2 and 10 min^−1^. (c) Phase space projection of the cyclin B / APC/C nullcline (in blue), cyclin B - Cdk1 nullcline (in orange), APC/C nullcline (in green) and oscillation (in gray) for the indicated values of *k*_*s*_. All other parameters are given in Table 1.

Finally, we studied the robustness of the cell cycle oscillator in more detail by exploring its existence when changing the properties of the two underlying switches. More specifically, we first studied the effect of the position of each switch by changing *C*_*x*_ and *C*_*xy*_, without changing *b*_*y*_ and *b*_*z*_ (determining their width). Figure 10(a) shows the resulting region in parameter space where oscillations were found. For comparison, we also show the oscillatory region when abolishing the bistability in either the (cyclin B, active Cdk1) switch [see Figure 10(b)] or the (active Cdk1, active APC/C) switch [see Figure 10(c)]. We find that oscillations persist over a wider range of parameters when both response curves are bistable (wider than the sum of both oscillatory regions in Figure 10(b) and (c)]. Another important observation is that the Cdk1 activation threshold *C*_*x*_ is required to be larger than the APC/C activation threshold *C*_*xy*_ to have oscillations. To gain a better understanding of how these switches interact to determine whether oscillations occur or not, we looked at the system dynamics in the presence of only a single underlying S-shaped response curve.

**Figure 10.**
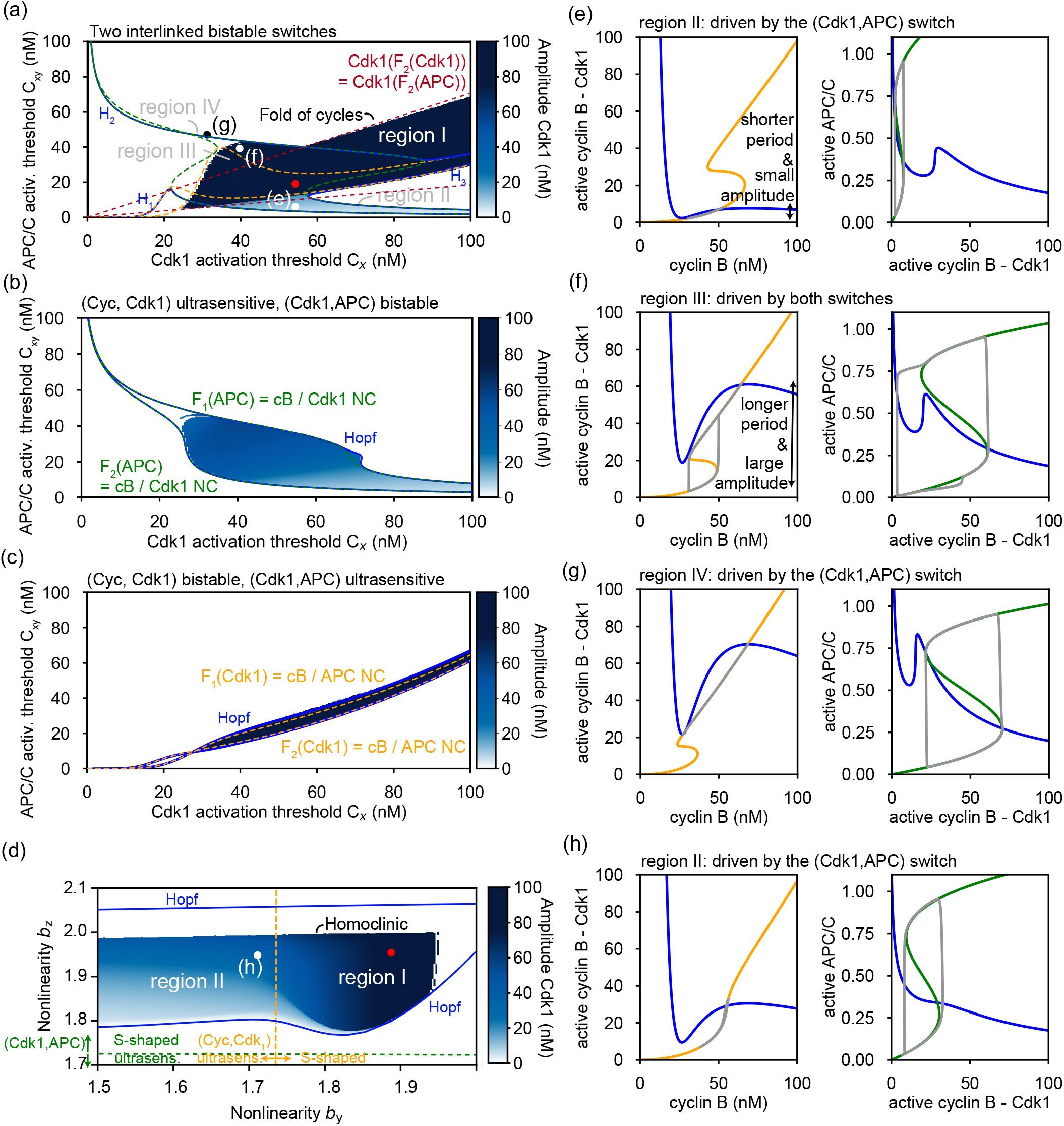
Phase diagrams of model (6)-(8) showing different oscillatory regimes. (a-c)(*C*_*x*_, *C*_*xy*_)-phase diagrams where the amplitude of the active Cdk1 oscillation is represented with the blue color map, the blue lines represent the Hopf bifurcations, the orange dashed lines correspond to the intersection of F_1,2_ (Cdk1) with the cyclin B nullcline and the green dashed lines represent the intersection of the F_1,2_ (Cyc) with the cyclin B - Cdk1 nullcline. The parameters are given in Table 1 for panel (a). The (cyclin B, active Cdk1) has been made ultrasensitive using *b*_*y*_ = 1.7 in (b). The (active Cdk1, APC/C) has been made ultrasensitive using *b*_*z*_ = 1.7 in (c). (d) shows the (*b*_*y*_, *b*_*z*_)-phase diagram for parameters given in Table 1. The amplitude of the active Cdk1 oscillation is represented with the blue color map, the blue curves represent a Hopf bifurcation, and the dashed orange and green lines represent the transition of (cyclin B, active Cdk1) and (active Cdk1, APC/C) switches from S-shaped to ultrasensitive respectively. (e,f,g,h) Phase space projection of the cyclin B / APC/C nullcline (in blue), cyclin B - Cdk1 nullcline (in orange), APC/C nullcline (in green) and oscillation (in gray). Parameter values are *C*_*x*_ = 55*nM*, *C*_*xy*_ = 5*nM* for (e), *C*_*x*_ = 40*nM*, *C*_*xy*_ = 40*nM* for (f), *C*_*x*_ = 30*nM*, *C*_*xy*_ = 46*nM* for (g), and *b*_*y*_ = 1.7, *b*_*z*_ = 1.95 for (h). The other parameters are given in Table 1.

When the cell cycle oscillator has only one underlying bistable switch, we found that the oscillations (again originating at a Hopf bifurcation) are well predicted by the condition that the projected nullclines cut each other in the middle branch of the S-shaped nullcline. These conditions are shown in green in Figure 10(b) and in orange in Figure 10(c). We then plotted these same conditions (when one nullcline crosses the folds of the S-shaped nullcline) when both underlying switches were bistable, see Figure 10(a). Moreover, we added the condition [Cdk1[F_2_(Cdk1)] = Cdk1[F_2_(APC)]] in red. In doing so, we found that these conditions nicely delineated the regions of oscillations, and we identified four different oscillatory regions. Region I corresponds to the oscillations that were driven by the (cyclin B, active Cdk1) switch as shown in 8. By decreasing the APC/C activation threshold C_*xy*_ the cyclin B / APC/C nullcline eventually crosses the bottom branch of the cyclin B - Cdk1 nullcline [when F_2_(Cdk1) = cB / APC NC, in dashed orange], see Figure 8(b). When this occurs, the oscillator changes its behavior as it becomes driven by the (active Cdk1, APC/C) switch, see Figure 10(e). Here, in region II in Figure 10(a), the oscillations are of low amplitude in cyclin B - Cdk1 activity. Figure 10(a) shows two more regions of oscillations, indicated as region III and IV. Region III occurs when the Cdk1 activity at the right fold of the APC/C nullcline coincides with that at the right fold of the cyclin B - Cdk1 nullcline [when Cdk1(F_2_(APC)) = Cdk1(F_2_(Cdk1))]. The oscillations are then driven in a more complex way by both bistable switches, see Figure 10(f). Cyclin B - Cdk1 is quickly activated while APC remains inactive due to Cyclin B - Cdk1 activity is below APC activating threshold. Thus, activity continues to increase following the nullcline until APC threshold is crossed and degradation suddenly increases. In region IV, the APC/C activation threshold surpasses F_1_(Cdk1) determined by *C*_*y*_ and the system remains in the upper branch of the nullcline (the active state) while the oscillations are driven by the (active Cdk1, APC) switch [Figure 10(g)]. Oscillations in region IV only exist in a very narrow region for low values of the Cdk1 activation threshold *C*_*x*_ and increasing values of the APC/C activation threshold *C*_*xy*_. The narrow shape of region IV is determined by the particular shape of the nullcline, which have a similar slope and cross each other relatively fast when activating thresholds are changed. Different values of *b*_*y*_, *b*_*z*_ lead to a wider region in parameter space. The four regions indicated in Figure 10 included oscillations which encircle the (active Cdk1, APC) switch alone or oscillations which encircle both switches simultaneously. It is, however, also possible to find oscillations which encircle only the (cyclin B, active Cdk1) switch with low amplitude APC/C oscillations. This particular regime is less likely to appear because higher APC/C-driven degradation rates (*d*_2_) are required. In other words, less degradation mediated by APC allows for low amplitude APC/C oscillation while fully degrading cyclin B. Again such oscillations provide additional robustness to parameter changes when the two switches are not simultaneously functional.

Next, we changed the width of the bistable switches by varying *b*_*y*_ and *b*_*z*_, while keeping their position fixed, see Figure 10(d). The first observation is that wider bistable switches (higher values of *b*_*y*_ and *b*_*z*_) favor oscillations driven by the (cyclin B, active Cdk1) switch, while also having a complete activation of APC/C (region I). Thus, the presence of the two bistable switches confers robustness by ensuring both cyclin B - Cdk1 and APC/C are fully activated and deactivated with high amplitude oscillations, which represents an advantage over a single switch. The oscillations persist even when the (cyclin B, active Cdk1) switch becomes ultrasensitive (region II). Instead, they are now driven by the (Cdk1, APC/C) switch: the limit cycle encircles it closely, see Figure 10(h). While the APC/C activity is still fully (de)activated, the oscillations in Cdk1 activity are of lower amplitude. These simulations show that both built-in switches can serve as drivers of cell cycle oscillations, building in robustness to parameter changes. The oscillatory regime is bounded between two Hopf bifurcations. As one increases *b*_*z*_ the system starts to oscillate at the first Hopf bifurcation. When increasing *b*_*z*_ further, the oscillation period and the time spent in a high activity state increases. Such an increase in the period is the result of approaching a homoclinic bifurcation that occurs before the second Hopf bifurcation is encountered.

## Discussion

Upon fertilization, the early *Xenopus laevis* frog egg quickly divides about ten times to go from a single cell with a diameter of a millimeter to several thousands of cells of somatic cell size. These early cell divisions occur very fast and largely in the absence of transcriptional regulation. They are driven by a cell cycle oscillator with periodic changes in the activity of the enzymatic protein complex cyclin B with cyclin-dependent kinase 1 (Cdk1). This cell cycle oscillator is driven by negative feedback involving 3 key components: active cyclin B - Cdk1 activates the Anaphase-Promoting Complex – Cyclosome (APC/C), which triggers the degradation of cyclin B Yang and Ferrell (2013); Rombouts et al. (2018). In the absence of this negative feedback, Cdk1 activity was found to respond in a bistable manner to changes in cyclin B concentration Pomerening et al. (2003); Sha et al. (2003) [see Figure 1(a)-(c)]. As a result, the cell cycle oscillator is an example of a typical *relaxation oscillator*, which has been shown to be important to generate robust, large-amplitude oscillations with tunable frequency Brandman et al. (2005); Tsai et al. (2008). Without a built-in bistable switch, changes in the amplitude would strongly correlate with changes in the frequency.

Interestingly, recent studies have shown that a second bistable switch exists in the system, as APC/C gets activated in a bistable manner by active Cdk1 Mochida et al. (2016); Rata et al. (2018); Kamenz et al. (2021) [see Figure 1(b)]. Motivated by these findings, we investigated here how a cell cycle oscillator built on two bistable switches would work. Over the years, many cell cycle oscillator models have been built, some focusing more on the detailed components and interactions in the regulatory network, while others focused more on the dynamical design principles required to make a biochemical circuit oscillate (for a review, see Novák and Tyson (2008); Ferrell et al. (2011)). In this work, we introduced the equations describing the temporal evolution of cyclin B concentration, the fraction of active Cdk1 and APC/C activity based on phenomenological response curves approximating experimental measurements. Using this modular approach to reduce the complexity, we then focused on understanding the dynamics of this cell cycle control system using theory of bifurcations in dynamical systems (for a review on such a dynamical approach in molecular cell biology, see Tyson and Novak (2020)).

The basic dynamics is most easily understood focusing on a cell cycle oscillator based on one switch, such as the ones described by Eqs. (13)-(14) and (15)-(16). The system lives in between the two states defined by the underlying bistable switch. Previous work illustrated that there exists a critical parameter, the ratio between synthesis and degradation rates *k*_*s*_*/d*_2_*C*_*z*_, that determines the system dynamics Rombouts et al. (2018). When this ratio is small (degradation dominates), the system finds itself in a stationary state of low activity on the lower branch of the S-shaped nullcline (inactive state). Instead, when the ratio is sufficiently high (synthesis dominates), a stationary state of high activity on the upper branch of the S-shaped nullcline (active state) results. In between, synthesis is fast enough to activate the system, but slow enough to allow degradation in the active state to reset the system to its inactive state. As a result, the system toggles between the two states producing a periodic oscillation (oscillatory state). We analyzed these dynamics in detail for the cell cycle oscillator built on a (cyclin B, active Cdk1) switch (13)-(14), and one built on a (active Cdk1, APC) switch (15)-(16).

When the two bistable switches are combined the resulting dynamics is more complex as the system can potentially transition between multiple different combinations of the previously described states. The system may find itself in a regime were both switches are either in the active or the inactive state, leading to a stationary state. However, there are also multiple ways in which the system can oscillate. On the one hand, one single switch can drive the oscillations by itself (encircling its S-shaped curve), while the system remains on the upper or lower branch of the S-shaped curve of the second switch as can be seen in Fig. 10(e) and (g). On the other hand, we also found that the two switches can work together to drive the oscillation under suitable conditions. The main factor contributing to this coordination is the positioning of the switches in the space (cyclin B, active Cdk1, active APC/C) [or equivalently: (*X, Y*, *Z*)], which is essentially controlled by the activation thresholds (*C*_*x*_, *C*_*xy*_) and the scale of the response curve (*C*_*y*_, *C*_*z*_) (see Fig. 10). The ability to generate oscillations with just a single switch confers robustness to the system by allowing oscillations to rely on the (cyclin B, active Cdk1) switch when the (active Cdk1, APC) switch is not functioning, and the other way around, thus extending the region of oscillations in parameter space. Indeed, measurements have shown that fast cell cycle oscillations persist after the first cell cycle despite the fact that (cyclin B, active Cdk1) bistability is absent Tsai et al. (2014). Recent work on circadian clocks and NF-κB oscillators driven by a transcriptional negative feedback loop also showed that combinations of multiple (potentially redundant) repression mechanisms are used to generate strong oscillations (sharper responses) Jeong et al. (2022). Our results are consistent with those findings and they are also in line with our previous work on combining multiple different functional motifs De Boeck et al. (2021). Here, by parametrizing bistable response curves based on experimental measurements, we extended our analysis of the dynamics characterizing the existence of the different oscillatory regimes related to the main bifurcations.

The distinction between the active and inactive states becomes less clear when the width of the bistable switch vanishes leading to an ultrasensitive, more graded response. Assuming the parameters used are the most representative of the experimental response curves, one may conclude the oscillations rely more on the (active Cdk1, APC) switch than the (cyclin B, active Cdk1) switch since the oscillations persist in the absence of bistability in (cyclin B, active Cdk1), while the reverse is not true. However, the amplitude of the oscillation reveals that both switches are required to be bistable in order to have oscillations with a complete activation of cyclin B - Cdk1 and APC/C.

Multiple questions arise regarding the influence of two interlinked bistable switches when more realistic features of the cell are considered. Frog egg extract experiments in a spatially-extended context have led to the observation that waves of mitosis are able to coordinate mitotic entry Chang and Ferrell Jr (2013); Gelens et al. (2014); Afanzar et al. (2020); Nolet et al. (2020), which was attributed to the presence of the (cyclin B, active Cdk1) switch. Such traveling biochemical waves can synchronize the whole medium and ensure that cell division is properly coordinated in the large frog egg. However, such spatial waves have typically been studied using simple cell cycle models relying on a single switch and are described by two variables Chang and Ferrell Jr (2013); Gelens et al. (2014); Nolet et al. (2020). The influence of multiple bistable switches on mitotic waves remains to be investigated Chang and Ferrell Jr (2013); Afanzar et al. (2020); Nolet et al. (2020). The ability to gradually increase (decrease) the presence of the (cyclin B, active Cdk1) switch by *b*_*y*_, as well as control the time-scale separation by *ϵ*_*y*_ and *ϵ*_*z*_, could also be interesting to study transitions from trigger waves to sweep waves Vergassola et al. (2018); Hayden et al. (2022); Di Talia and Vergassola (2022).

Another consequence of the presence of two interlinked bistable switches is the appearance of two types of excitable regimes. Type I excitability appears when the period diverges close to the bifurcations involved in the creation or destruction of the limit cycle (i.e., or the limit cycle emerges with zero frequency). We talk about excitability of Type II when the period does not diverge and remains almost constant when approaching the bifurcation. In this last case, the limit cycle appears with a finite frequency. Type I excitable behavior is expected to arise as a result of coexistence of different equilibria together with oscillatory dynamics, and is related with two main bifurcations: the SNIC, where the limit-cycle collides with a saddle-node bifurcation, and with homoclinic bifurcations Izhikevich (2007); Moreno-Spiegelberg et al. (2022). The type I excitability reported here is related with the second case. Type II excitability is mediated by the presence of a fold of cycles together with a subcritical Hopf bifurcation, or by a supercritical Hopf bifurcation with a Canard explosion, resulting from timescale separation or relaxation-like character of the bistable switches Meron (1992). The different excitable regimes may lead to different excitable excursions with different features in phase space and their associated spatial dynamics, such as traveling pulses.

Describing cell cycle oscillations using two bistable switches can also be enlightening under nuclear compartmentalization. Recently, *in vitro* reconstitution of nuclear-cytoplasmic compartmentalization showed how the nucleus influences the trajectory in phase space encircling (cyclin B, active Cdk1) switch Maryu and Yang (2022), previously also studied in Pomerening et al. (2005). Nuclear compartmentalization can lead to changes of the underlying bistable response curves Rombouts and Gelens (2021) or incorporate spatial positive feedback due to phosphorylation promoting nuclear translocation Santos et al. (2012); Chae et al. (2022), which can be interpreted as an additional bistable switch. The nucleus has recently also been shown to serve as a pacemaker to drive mitotic waves Nolet et al. (2020); Afanzar et al. (2020). As the properties of such pacemaker-driven waves depend on the oscillator properties Rombouts and Gelens (2020), it would be also relevant to study the effect on such waves when having an oscillator built on multiple switches.

Activation and deactivation thresholds are also susceptible to changes regulated by other processes. The shape of response curve and the associated threshold can be strongly affected by changes in protein concentration due to multiple factors within the cell, such as DNA damage Kwon et al. (2017); Stallaert et al. (2019), the relative amount between Cdc25 and Wee1 Tsai et al. (2014), or spatial differences in concentration Rombouts and Gelens (2021). During cell cycle progression cyclin B Cdk1 and APC/C activation (deactivation) thresholds may not change simultaneously, serving as a mechanism for (de)coupling different processes and adjusting the timing of different cell cycle events.

## Acknowledgements

P. P. -R acknowledges support from the European Union’s Horizon 2020 research and innovation programme under the Marie Sklodowska-Curie grant agreement no. 101023717. D.R-R. is supported by the Ministry of Universities through the “Pla de Recuperació, Transformació i Resilència” and by the EU (NextGenerationEU), together with the Universitat de les Illes Balears.

## Methods

### Numerical methods

Numerical integration of the different models has been performed using custom codes in python based on the ODE integrator Scipy.integrate.odeint. The numerical codes that were used are available through GitLab (https://gitlab.kuleuven.be/gelenslab/publications/cb-cdk1-apc-model). Numerical continuation of stationary solutions, periodic orbits, and bifurcations have been performed using the continuation software AUTO07p Doedel and Oldeman (2009).

